# Enhanced laterally resolved ToF-SIMS and AFM imaging of the electrically conductive structures in cable bacteria

**DOI:** 10.1101/2021.01.27.428206

**Authors:** Raghavendran Thiruvallur Eachambadi, Henricus T. S. Boschker, Alexis Franquet, Valentina Spampinato, Silvia Hidalgo-Martinez, Roland Valcke, Filip J. R. Meysman, Jean V. Manca

**Affiliations:** UHasselt – X-LAB, Agoralaan – Gebouw D, 3590 Diepenbeek, Belgium; Department of Biotechnology, Delft University of Technology, Van der Maasweg 9, 2629 HZ Delft, The Netherlands; Department of Biology, University of Antwerp, Universiteitsplein 1, 2610 Wilrijk, Belgium; Materials and Components Analysis – Compositional Analysis, Imec vzw, Kapeldreef 75, 3001 Leuven, Belgium; UHasselt – Molecular and Physical Plant Physiology, Agoralaan – Gebouw D, 3590 Diepenbeek, Belgium

## Abstract

Cable bacteria are electroactive bacteria that form a long, linear chain of ridged cylindrical cells. These filamentous bacteria perform centimeter-scale long-range electron transport through parallel, interconnected conductive pathways of which the detailed chemical and electrical properties are still unclear. Here, we combine ToF-SIMS (time of flight secondary ion mass spectrometry) and AFM (atomic force microscopy) to investigate the structure and composition of this naturally-occurring electrical network. The enhanced lateral resolution achieved allows differentiation between the cell body and the cell-cell junctions that contain a conspicuous cartwheel structure. Three ToF-SIMS modes were compared in the study of so-called fiber sheaths (i.e., the cell material that remains after removal of cytoplasm and membranes and which embeds the electrical network). Among these, fast imaging delayed extraction (FI-DE) was found to balance lateral and mass resolution, thus yielding multiple benefits in the study of structure-composition relations in cable bacteria: (i) it enables the separate study of the cell body and cell-cell junctions, (ii) by combining FI-DE with in-situ AFM, the depth of Ni-containing protein – key in the electrical transport – is determined with greater precision, and (iii) this combination prevents contamination, which is possible when using an ex-situ AFM. Our results imply that the interconnects in extracted fiber sheaths are either damaged during extraction, or that their composition is different from fibers, or both. From a more general analytical perspective, the proposed methodology of ToF-SIMS in FI-DE-mode combined with *in-situ* AFM holds great promise for studying the chemical structure of other biological systems.

---

Cable bacteria are multicellular microorganisms that form long un-branched filaments and belong to the *Desulfobulbaceae* family^1^. They are the focus of interdisciplinary research due to their unique capability of conducting electrical currents over centimeter distances^2,3^, a process also known as long-distance electron transport (LDET). Cable bacteria have been found to thrive in different environments such as fresh water^4,5^ and marine sediments^6,7^ and have also been found in different parts of the world^7^. Cable bacteria display a distinct morphology with parallel ridges running along the length of the filament^1,8,9^. Scanning Electron Microscopy of cable bacteria cross-sections revealed the presnce of fibers of about 50 nm in diameter under the ridges and a cartwheel structure at the junctions^8^. These fibers are embedded in the periplasm (i.e., in space between the cytoplasmic membrane and the bacterial outer membrane) and were suspected to be the conductive structures^8^ (Figure 1 A-C).

**Figure 1.**
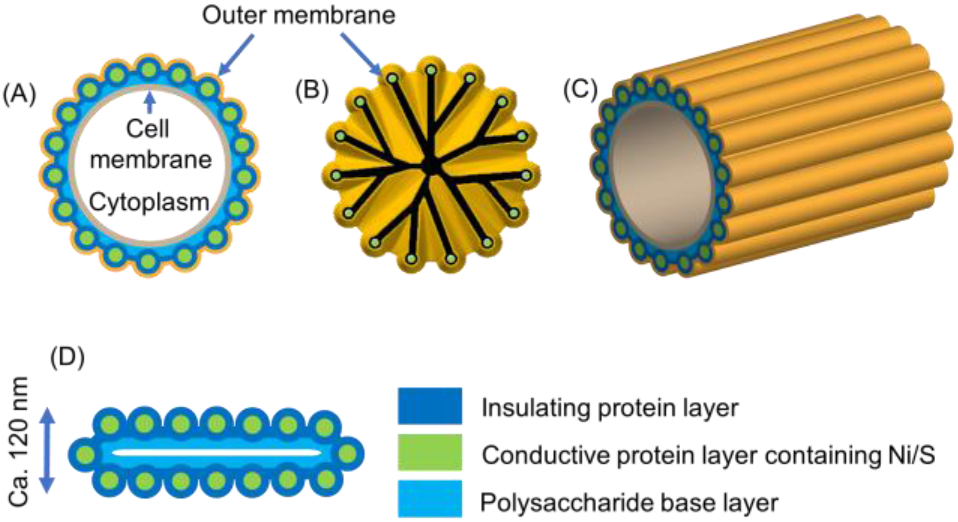
Schematic of a cable bacterium. Cross-section of a cable bac-terium filament at the center of a cell (A) and at the junction (B) showing the cartwheel structure. 3D representation of the ridged cell (C). After the sequential extraction procedure, a flat ca. 120 nm thick fiber sheath is obtained, seen in (D). The spokes of the cartwheel are shown in black as its composition is unknown (adapted from Cornelissen et al.8 and Boschker et al.11)

**Figure 2.**
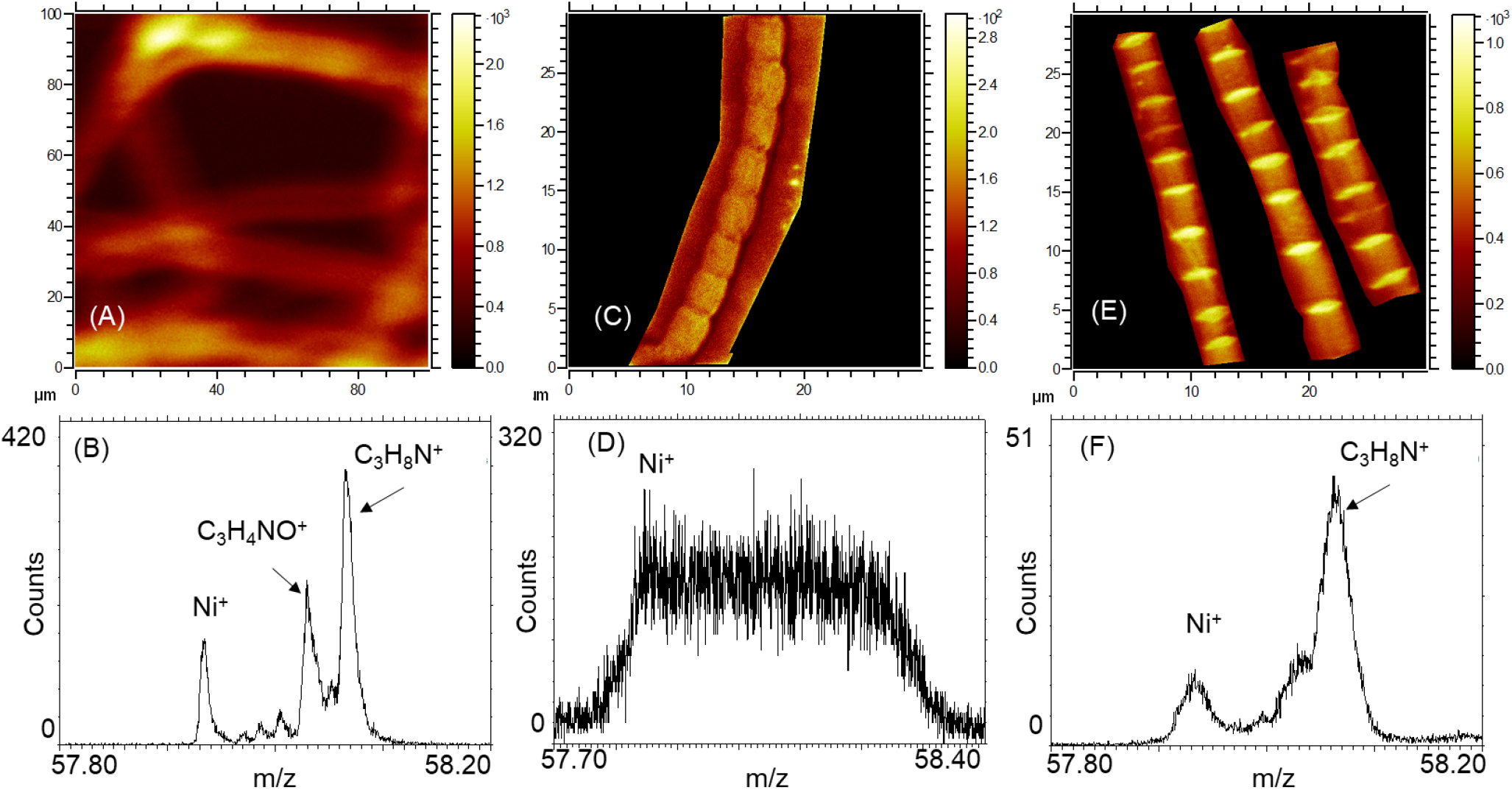
Study of ToF-SIMS imaging modes on the lateral and mass resolution: HCB (A and B), FI-DE (C and D) and FI mode (E and F). (A), (B) and (C) are the total intensity images. The lateral resolution of FI-DE (E) and FI (C) is better than HCB (A). On the other hand, the mass resolution (m/Δm) of Ni+ of HCB mode (B) was 7000, whereas in FI-DE mode (D) was lower at 2100, sufficient to resolve Ni+ signal. Mass resolution in FI mode is insufficient to resolve Ni+.

Recently, Meysman et al. experimentally investigated the conductivity of these fibers^9^. A sequential extraction procedure was developed^8^ (see experimental section), by which the fiber structures can be isolated from cable bacterium filaments^8^. After chemical removal of cytoplasm and membranes, a so-called fiber sheath remains, which embeds the periplasmic fibers^9–11^. The fiber sheath flattens when air-dried (Figure 1D), and the top part of this fiber sheath mirrors the bottom part due to its cylindrical symmetry^8^.

Meysman et al. demonstrated that fibers sheaths were indeed highly conductive^9^. Fiber sheaths were placed on top of two gold pads with a non-conductive oxide spacing, and when applying a potential difference between the two pads, a flow of current indicated that the periplasmic fibers are the conductive conduits. These resuls were subsequently confirmed with conductive atomic force microscopy (C-AFM)^[10]^. The conductivity of these periplasmic fibers (> 20 S/cm) not only rivals that of doped organic semiconductors^9^, but the length scale of electron transport is also more than three orders of magnitude longer than previously known for microbial structures^3^. Therefore, there is a strong interest in this material for future biodegradable electronic applications.

In order to better understand the unique electrical properties of cable bacteria, a key challenge is to unravel the composition of the electrically conductive fibers. Here, we combine ToF-SIMS and AFM to investigate the structure and composition of this natural-occurring electrical network of fibers with enhanced lateral resolution. Mass spectrometry-based chemical imaging is widely used for different types of cellular analyses^12–14^. Various SIMS techniques are available, of which ToF-SIMS and nanoscale secondary ion mass spectrometry (NanoSIMS) are routinely utilized. Of the two, NanoSIMS provides the best lateral resolution (< 50 nm), but is limited in the number of elemental masses that can be simultaneously detected^15–19^. With isotopic labelling, NanoSIMS can be used to localize the incorporation of different elements (C, N, S) within the cell compartments^20–22^, and this technique has also been recently applied to cable bacteria to investigate the relation between LDET metabolism and filament growth^20–22^.

ToF-SIMS uses a polyatomic or gas cluster ion source in addition to monoatomic sources, and as such, it is less destructive as compared to NanoSIMS. Bi_3_ ^+^ ion source was found to be both surface-sensitive as well as providing the best imaging contrast^23^. Large molecular fragments up to 1000 Da can be analyzed, and with a typical lateral resolution of 100 nm – 10 μm^24^. Generally, two different modes are employed in ToF-SIMS^25^. The first one is the high current bunched (HCB) mode, also referred to as mass-spectrometry mode, which targets high mass resolution but has a restricted 2-10 µm lateral resolution^25,26^. Mass resolution can be defined as the ability to distinguish two peaks of slightly different mass-to-charge ratios (m/Δm) in a mass spectrum. HCB mode uses three electrostatic lenses and a primary ion buncher system, ensuring short pulses of less than one ns. Such short pulse duration results in a typically high mass resolution m/Δm > 10,000. The second ToF-SIMS mode is the burst alignment mode or fast-imaging (FI) mode, where a high lateral resolution of about 400 nm is obtained but with a loss of mass resolution^27^. A narrow beam is used in FI mode, with a beam diameter well below one micron using two electrostatic lenses. The time width of the primary ion pulse is in the order of tens of nanoseconds, leading to a low unit mass resolution (m/Δm ∼ 200)^25–27^. Mass resolution can be improved by maintaining the lateral resolution of FI mode using delayed extraction. This third ToF-SIMS mode is termed as fast imaging delayed extraction (FI-DE). In delayed extraction, ion extraction is decoupled from ion generation by switching off the extraction voltage for several nanoseconds after firing the primary ion pulse. A plume of ions is obtained just over the surface, a field-free emission of secondary particles. Due to this decoupling, the long primary ion pulse required to obtain a high lateral resolution does not affect the mass resolution^25,27^. The plume moves away from the surface before the extraction voltage is switched on, because of which the topographic effects are reduced, the number of secondary ions collected increased, and sharper lateral images with a better signal are obtained. Quite recently, Benettoni et al. were able to obtain a high lateral resolution of ∼100 nm on a chessboard sample, with a mass resolution in the order of 5000. However, the lateral resolution was reduced to 222 nm on an algal biofilm^28^.

Recently, TOF-SIMS analysis combined with in-situ AFM has generated the first insights into the conductive network of cable bacteria^11^ (see Figure 1D for a schematic representation of results). To this end, ToF-SIMS HCB analysis using Bi_3_ ^+^ was combined with interlaced argon cluster sputtering and applied to fiber sheaths. This provided high-resolution depth profiles of both organic and inorganic constituents at low lateral resolution^11^. High surface counts were recorded for amino acid fragments, including aromatic amino acids in both positive and negative modes. Nickel and Sulphur signals showed subsurface peaks in the positive and negative mode, suggesting that the fiber’s central core is protein-rich with Ni and S. After 150s of sputtering, signals from the oxygen-rich fragments, including carbohydrate specific ions peaked while the signals from nitrogen-containing fragments levelled off. By combining results from other complementary characterization techniques, a structural model of the fiber sheath was made (Figure 1D): fibers are made of protein, lying on top of a polysaccharide-rich base layer, most likely consisting of peptidoglycan. The fiber itself is made of a Ni-rich protein core surrounded by a thin layer of Ni-deficient protein, which is termed as a fiber core/shell structure^11^. Although the HCB mode was instrumental in identifying various fragments with a high mass resolution, it comes with a sacrifice of lateral resolution that does not allow to separately study the composition of the fibers and the cartwheel structure at the junctions^11^.

To provide more detail on the conductive fibers present in the fiber sheath, we employed and compared the three mentioned ToF-SIMS modes. The FI-DE mode, which balances lateral and mass resolution, in combination with in-situ AFM is expected to offer the unique benefit of a direct and more detailed depth calibration.

## EXPERIMENTAL SECTION

### Sample preparation

Sediments containing cable bacteria were collected from a salt marsh creek bed. These sediments were sieved, homogenized, repacked in PVC core liner tubes (diameter 40 mm), and were subsequently placed in aerated, artificial seawater. These incubations are known to consistently develop thick, ca. 4 μm diameter, cable bacterium fialments, which facilitates their isolation from the sediment and fiber sheath extraction.

To collect the cable bacterium filaments and extract the fiber sheaths, a small amount of sediment was placed on a microscope cover slip. Multiple 20 μL droplets of Milli-Q water were placed near the sediments. Under a stereomicroscope, filaments were picked from the sediments using custom-made glass hooks made from Pasteur pipets. Filaments were cleaned and washed at least six times by transferring them between droplets. The cleaned intact filaments were subsequently incubated in a 20 μL droplet of sodium dodecyl sulfate (SDS) for 10 minutes, followed by six MilliQ droplet washes. Filaments were further subjected to a 10-minute incubation in a 20 μL of 1mM sodium ethylenediaminetetraacetate (EDTA) solution, again followed by six washes in Milli-Q^8^. The extracted material represents the fiber sheath containing the conductive fibers.

Fibers sheaths were deposited as clumps on a 1 cm x 1 cm diced Au covered Si wafer for ToF-SIMS analysis. Samples were first imaged in an optical microscope to identify areas to be analyzed. Three samples prepared on different occasions were analyzed under HCB and FI-DE mode, and five areas from two clumps of fiber sheaths prepared during the same run were analyzed under combined ToF-SIMS/AFM mode.

For conductive AFM, one filament was deposited on a 1 cm x 1 cm diced SiO_2_ covered Si wafer (Figure S5). This wafer was then affixed onto a steel disc using silver paste (EM-Tec AG44 conductive silver paint). The detailed procedure can be found elsewhere^10^. Four replicates were analyzed using C-AFM.

### ToF-SIMS analysisToF-SIMS analysis

ToF-SIMS analysis was performed using TOF.SIMS NCS (IONTOF GmBH, Germany) located at imec, Leuven (Belgium). For HCB mode, ToF-SIMS was carried out in interlaced mode using Bi3^+^ analysis beam (30keV, current ∼0.35 pA, 100×100 μm^2^ area, 256×256 pixels) and Ar4000^+^ gas cluster ion beam (Ar GCIB, 10keV, current 1nA, 400×400 μm^2^ area). Fast imaging (FI) was done by finetuning the existing factory settings (Bi3^+^, 30 keV, ∼0.12 pA, 30 × 30 μm^2^, 512 x 512 pixels). Only the surface scans were obtained in the case of FI mode. For FI-DE, the existing setting for fast imaging was initially set with Bi3+ ions (30keV, current ∼0.15pA, 30×30 μm^2^ area). Parameters for delayed extraction were optimized, namely, delay time, the analyzer lens voltage, X/Y analyzer deflection plates, and surface and virtual drift potential (VDP)^27^. A delayed extraction of 85 ns was found to be appropriate. A cycle time of 50 μs was used. In both cases, filaments were not necessarily sputtered until only the substrate remained, as shown in Figure 4.

**Figure 3.**
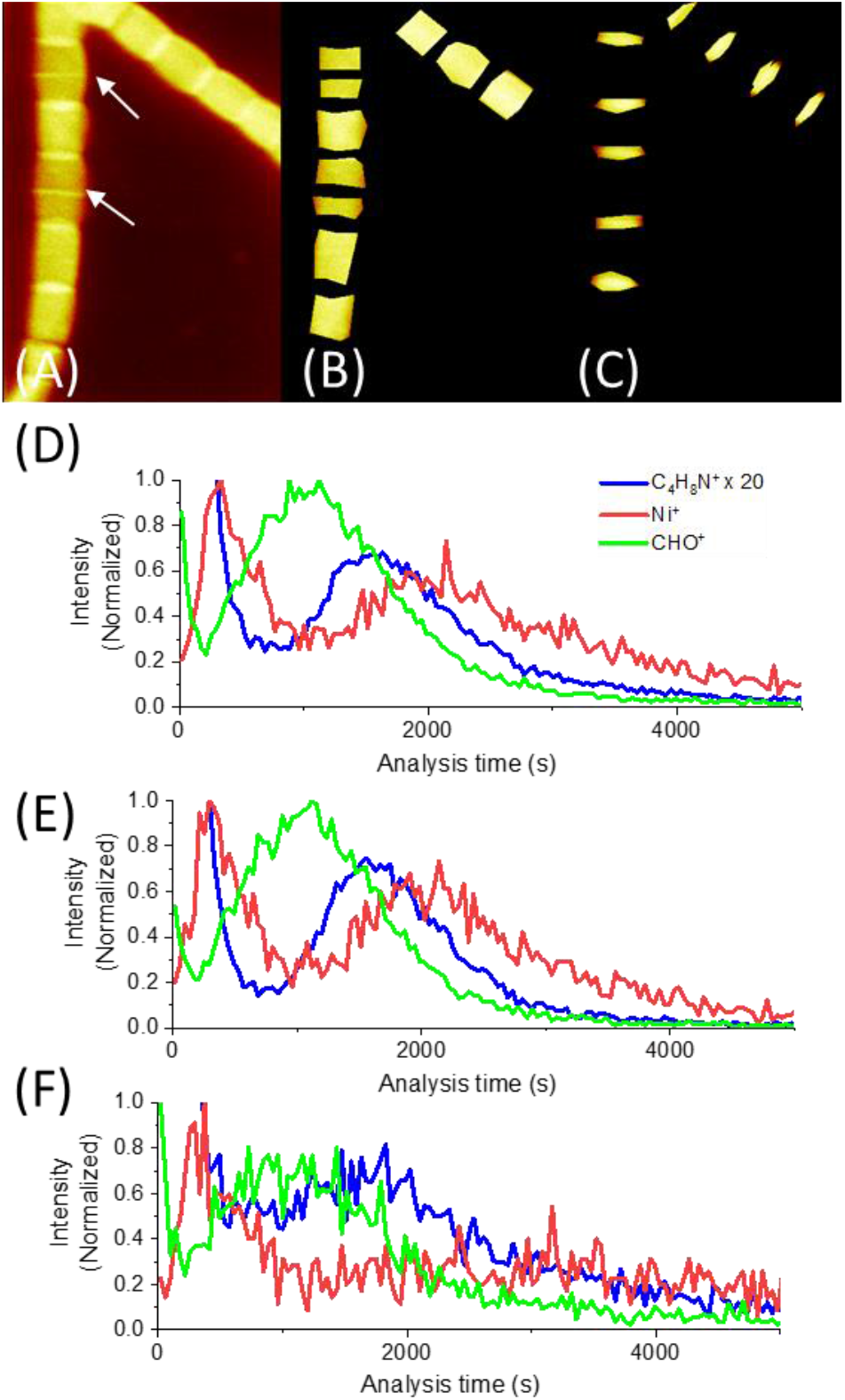
(A) Lateral total intensity image of two cable bacteria filaments, with two white arrows showing the newly forming cell junctions. The enhanced lateral resolution of FI-DE enabled the study of signals from various fragments from the body (B) and the junctions (C) separately. Trends of C_4_H_8_N^+^, Ni^+^, and CHO^+^ signals, normalized to their highest intensity, from the filaments (D), cell bodies (E), and cell junctions (F). C_4_H_8_N^+^ signal was magnified by 20x in order to show the second peak in the profile.

**Figure 4.**
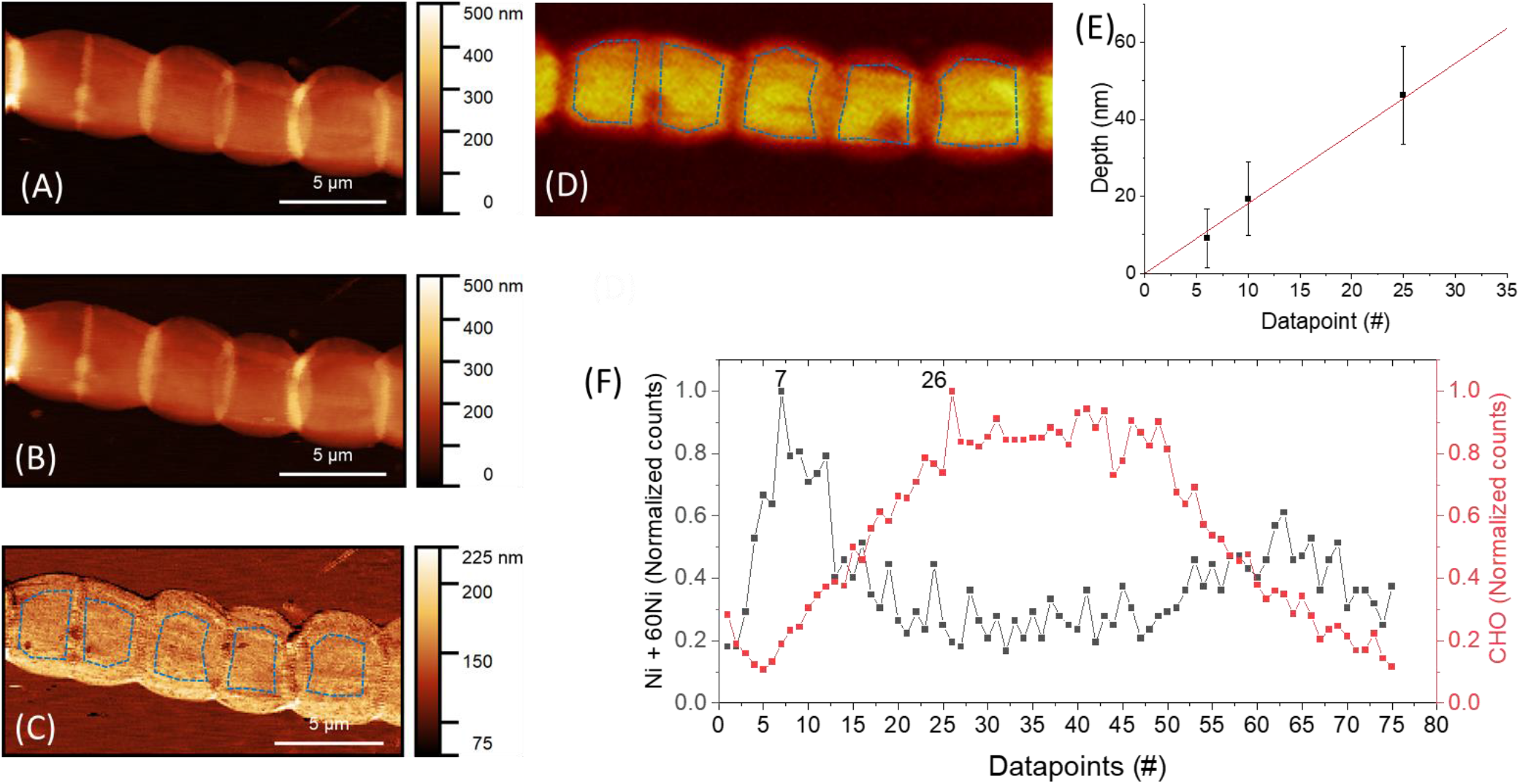
Combined ToF-SIMS/AFM analysis of a fiber sheath filament. An AFM image was captured (A) before sputtering and (B) after sputtering until datapoint 6. The total intensity ToF-SIMS image is shown in (D). (B) and (A) were aligned, and (B) was subtracted from (A), resulting in a difference image (C). Removed thickness was calculated by subtracting the substrate’s mean height from the mean height of the bacteria body (dotted blue polygons), plotted in (E). (F) shows the change of Ni+ and CHO+ counts (normalized, obtained from cell bodies, marked as dotted blue polygons in D) as a function of captured data points. Ni+ peak is seen at datapoint 7, and CHO+ signal reaches a maximum at datapoint 26. The second Ni peak is seen at datapoint 64.

**Figure 5.**
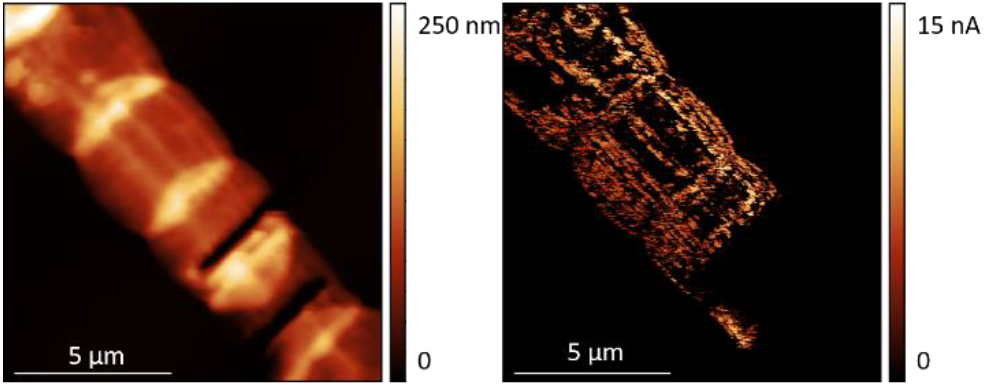
Figure 5. Conductive AFM image of a fiber sheath. (A) Topography map and (B) Current map. The fiber sheath is biased using carbon paste on the top-left corner (not seen). As the probe scans the surface from the bottom, structures still electrically connected to the carbon paste conduct current, which is visualized in (B).

SurfaceLab software (v7, IONTOF, Germany) was used for data analysis. In HCB mode, mass spectra were internally calibrated using C_2_H_3_ ^+^, C_3_H_4_ ^+^, C_3_H_5_^+^, C_4_H_5_, and Au^+^. Mass peaks were identified based on earlier ToF-SIMS work with cabel bacteria^11^. In FI-DE mode, mass spectra were internally calibrated using C_2_H_3_^+^, C_3_H_4_ ^+^, C_3_H_4_^+^, and C_4_H_5_^+^ .When necessary, lateral shift correction was done by using the *Shift correction* sub-program of the *Images* program. Regions of interest (ROIs) were created to analyze cell bodies and junctions separately (Figure 4). The peak list obtained from HCB mode was truncated to remove non-resolvable signals. Identified peaks from both HCB and FI-DE modes are provided in Tables S2 to S6.

Principal component analysis (PCA) was performed using OriginLab Pro v2020’s principal component analysis app. Three replicates of profiles of various fragments from HCB and FI-DE modes were separately analyzed after mass calibration with the same peak lists. The peak width of each signal was adjusted by overlapping the three spectra and fixing the peak width. Depth (or analysis time) profiles of selected peaks were imported into OriginLab Pro software, in which the profiles of individual fragments were normalized to their maximum value. Profile data of the various masses until the maxima of the oxygen-containing organic ions were used for PCA, identical to those of Boschker et al.^11^. Fragments used in PCA are indicated in Tables S2 to S6.

### Combined ToF-SIMS/AFM analysis

In-situ AFM was used in contact mode to analyze the depth at which the Ni signal maximum was found and to estimate the midpoint of the carbohydrate layer. PPP-EFM probes with a nominal spring constant of 3 N/m were used in contact mode. The AFM probe and the area to analyze were first aligned using a test area close to the area of interest. After ToF-SIMS obtained a scan of a known area, the coordinates were noted down by the Surfacelab software. Then, the sample was driven to the AFM part of the instrument. The area of interest was retrieved in the AFM by trial and error, and the Surfacelab software noted the coordinates of this area. The software calculates the lateral vector shift based on the two coordinates.

A 10 x 20 µm^2^ AFM image of the area to be sputtered was captured with a pixel size of about 78 nm. An enlarged area of about 15 x 30 µm^2^ was sputtered by Bi^3+^ ions with the conditions mentioned earlier. Sputtering was paused before and after the peak in Ni^+^ ion signal emerged. During the pause, the stage was moved within the same hybrid ToF-SIMS/AFM instrumental setup - without exposure to the lab atmosphere and therefore no contamination - to the in-situ AFM location. AFM imaging was taken from the sputtered area. These two images were leveled by mean plane subtraction, aligned using the “mutual crop” module of the *Gwyddion* software, and then the second image was subtracted from the first. In this difference image, the amount of material removed from the cell body can be determined. Another AFM image was taken at a point in the carbohydrate region.

### Conductive AFM

AFM analysis was done on a Multimode 8 (Bruker, Santa Clara, CA, USA) with Nanoscope V controller located at UHasselt. A CDT-NCLR probe with a nominal spring constant of 72 N/m was used. A fiber sheath was placed on a silicon substrate with a 100 nm thick SiO_2_ layer acting as an insulator. This substrate was then connected to a steel disc using silver paste (Figure S6). One end of the fiber sheath was electrically connected to the substrate holder, while the other end of the filament was left free. Bias is applied to the sample via the sample holder, and the conductive probe is electrically connected to the TUNA application module, which contains a current amplifier. This application module is, in turn, connected to the AFM controller. Current can only flow if there is an electrical connection between the substrate holder and the AFM probe, thereby completing the electrical circuit. Measurement was initially carried out in Scanasyst mode to obtain topography. After an area of interest was localized, AFM was switched to C-AFM mode, which works in contact mode.

Due to the high spring constant of the cantilever, specific areas from a cell were relatively easily removed by scratching a given area continuously in contact mode with a high force (15 µN) to disrupt more than half of the electrical connections within a cell. This creates a trench that goes all the way to the substrate. However, a much lower force (2.2 µN) was applied to gently scrape bacteria’s surface to visualize the electrical pathways, identical to our earlier work^10^.

## RESULTS AND DISCUSSION

### Comparison of lateral imaging and mass resolution

To qualitatively appreciate the mass resolution and quantitatively measure mass resolution, fiber sheaths were imaged using three different ToF-SIMS modes. Figure 2 shows cable bacterium filaments from a single sample preparation imaged with HCB (A and B), FI (C and D), and FI-DE (E and F). This illustrates the capabilities of these modes in terms of lateral imaging resolution (upper panels: A, C, and E) and mass resolution of ionized fragments (bottom panels: B, D, and F).

The HCB mode provides a high mass resolution spectrum (Figure 2A, B, S1). A high mass resolution of Ni^+^ (m/z = 57.93) of 7,000 was measured. Figure 2A clearly shows the poor lateral imaging resolution, in which the filaments appear fuzzy and much thicker than their nominal width of 4 to 5 µm. Using FI-mode led to a substantial improvement in lateral imaging resolution, with the cell junctions being resolved from the cell bodies (Figure 2C). However, the mass resolution was at best in the order of a few hundreds. For instance, the Ni^+^ signal was not resolved, because the low mass resolution (131), it also encompassed signals from C_2_H_4_NO^+^ (m/z = 58.029) and C_3_H_8_N^+^ (m/z = 58.065) (Figure 2C, D). This lack of mass resolution is insufficient for biological analysis.

In FI-DE mode, lateral resolution was sufficient to separate cell junctions from cell areas (Figure 2E), and the mass resolution of Ni^+^ was 2100 (Figure 2F). Complete spectra obtained from FI-DE can be seen in figure S1. Although the mass resolution does not match up to HCB, it was sufficient to resolve the Ni^+^ signal. FI-DE improved the lateral resolution, where the cell junctions and bodies are resolved. The mass resolution was found to be sufficient to resolve Ni and fragments of amino acids, polysaccharides, and others (see tables S2 – S6).

### Comparison of depth resolution

We compared the depth resolution between HCB and FI-DE. HCB uses a dual-beam, where Argon GCIB is used to sputter away the analyzed area. However, FI-DE is a single beam measurement where Bi_3_ ^+^ beam is also responsible for sputtering. Here we show that FI-DE has a better depth resolution compared to HCB.

Figure S2 shows the normalized three main trends in the sputtering-time depth profiles as found in HCB mode, based on the study by Boschker et al.^11^. The first one or two data points are usually related to surface transients, i.e., C- or N-based ions derived from contamination (this surficial zone extends until the local minimum of CHO^+^ signal). Below this, the first signal observed are high levels of nitrogen-containing fragments such as C_4_H_8_N^+^, a fragment of proline, an amino acid used in the synthesis of proteins, that stands for the profile of all amino acid fragments. Ni and its isotopes show a subsequent peak, and a third and broader peak is displayed by oxygen-containing fragments such as CHO^+^, most likely derived from carbohydrates in the peptidoglycan layer. The C_4_H_8_N^+^ shows a very high surface signal, indicating the presence of protein layer at the surface. The sub-surface peak in Ni has been linked to a Ni-containing protein that likely plays a role in the electron transport within the conductive fibers^11^. As the intensity of Ni^+^ decreases, oxygen-containing fragments such as CHO^+^ becomes higher in intensity. A second Ni peak is seen at about 164s of sputtering time. Although not entirely resolved, this second peak is likely due to the Ni-protein layer in the bottom part of the fiber sheath, which is essentially a mirrored duplicate of the part of the sheath away from the substrate.

Principal component analysis (PCA) of selected peaks (Figure S3A, C) showed clustering of oxygen-containing fragments and nitrogen-containing fragments with Ni. Boschker et al. proposed that the fiber sheath is made of a thin protein layer containing Ni-containing proteins and a polysaccharide-rich layer present under the protein layer^11^. However, in many cases, depth smearing occurs due to a possible different structure (cell bodies vs. junction). When cells retain an amount of cytoplasm after incomplete extraction, only the first subsurface Ni peak can be detected^11^. So an additional benefit of FI-DE as employed here is that it also enables the imaging of the bottom part of the fiber sheath.

A significant improvement made with FI-DE is that regions of interest (ROIs) can be separately defined for the cell body and the cell junction (Figure 3A-C). Signals from the body of cable bacteria this can be studied separately from the cell junctions, which reduces depth smearing, as junctions are thicker than the rest of the filament^8^. Cell junctions can be distinguished from the rest of the bacteria in FI-DE by the total counts measured at the junction, possibly due to the higher material yield. Cell junctions contain more material than the cell body, which implies more counts from this region (Figure 3A). Newly forming division planes, which are rings consisting of protein FtsZ^29^ were not considered in the junctions analysis since it is not known whether their structure is similar to that of an established division plane. Depth resolution was further improved in FI-DE imaging since more data points were obtained for a given thickness of material sputtered thanks to a single beam. The first seven data points, corresponding to the local minimum in CHO^+^ signal, are again related to the surface transient, i.e., organic contamination on the surface.

The filament signals in FI-DE mode consistently revealed a sharp sub-surface Ni peak and a distinctive second peak (Figure 3D). C_4_ H_8_ N^+^ has an intense surface signal seen earlier in HCB mode and a secondary peak just before Ni signal reaches a peak, which was not seen in HCB mode. The carbohydrate oxygen-containing fragment CHO^+^ is prominently present between the two peaks of Ni^+^ signals and nitrogen-containing signals. The second peak of Ni signal and the peak of C_4_ H_8_ N+ signal confirm that Ni is found in the protein, and the basal sheath is held together by a carbohydrate-containing layer. The second Ni^+^ peak and C_4_ H_8_ N^+^ peak from the cell bodies (Figure 3E) were sharper than the signals from the filaments. This is because junctions are thicker and possibly have a different composition due to the presence of the cartwheel structure. PCA analysis of the various identified fragments from cell bodies (Figure S3 B, D) showed a clustering of the nitrogen-containing fragments and the oxygen-containing fragments, similar to that seen in HCB mode.

### Combined ToF-SIMS/AFM study of fiber sheaths

Depth profiles in ToF-SIMS provide the intensity of various fragments as a function of time. By using in-situ AFM at appropriate intervals, analysis time can be translated in terms of distance. Depth patterns of various fragments as obtained by combining AFM and HCB mode were previously shown by Boschker et al.^11^. Here we combine AFM with FI-DE mode. Figure 4 describes how the distance of the Ni^+^ signal from the top surface is measured. Figure 4A is the height image before analysis commenced. The change of normalized intensity of the sum of two Ni isotopes, ^58^Ni^+^ and ^60^Ni^+^ and CHO^+^ signals during sputtering are given as a function of data points (Figure 4F). One data point corresponds to one scan by the Bi^3+^ ions of the given area. Profiles of ^58^Ni^+^ and ^60^Ni^+^ and CHO^+^ fragments in Figure 4F were obtained from the dotted blue polygons in Figure 4D. As the combined Ni signal went higher in intensity, the analysis was paused at datapoint 6, and another AFM image was taken (Figure 4B).

Subtracting Figure 4B from 4A gave a difference image, Figure 4C. Sputtered depth was measured by averaging the area within the dotted blue polygons of Figure 4C and subtracting from the substrate’s height. This corresponds to 9.1 ± 7.7 nm. Another AFM image was taken at data point 10 after the peak of Ni^+^ signal was crossed. A third AFM image was taken at datapoint 25, in the carbohydrate-rich region. After measuring the sputtered depths for data points 10 and 25, a line fit was made. The initial condition was that no material is removed before the commencement of analysis (i.e., y (x =0) = 0 nm). The peak of Ni^+^ signal was seen at datapoint 7, corresponding to 12.7 nm. Based on an average of five replicates, an average depth of 11.8 ± 0.6 nm was measured for the Ni maximum, slightly lower than the previously reported value of 15 ± 3 nm^11^. However, the midpoint of the flat region of the oxygen-containing organics varied between samples (between 32 nm and 68 nm, see table S1). This is probably because different filaments containing varied amounst of cytoplasm, although undergoing the same extraction procedure.

By combining in-situ AFM measurements with ToF-SIMS, we measured the depth of Ni^+^ without having to remove the sample out of vacuum. This is of importance to materials that are sensitive to exposure to atmospheric exposure. Also, keeping the sample within the equipment ensures the same instrument parameters such as vacuum and analysis beam conditions before and after AFM measurements. Lateral resolution obtained by FI-DE ensured a good correlation between SIMS and AFM data, and a more accurate determination of the depth at which Ni signals reached a peak in its counts.

### Composition of the cell junctions

Cell junctions contain the interconnecting structures that provide cable bacteria filaments with a redundant failsafe electrical network^10^. The cartwheel structure present at the junctions is suspected of containing the interconnecting structure, although there is no direct proof available^8,10^. Interestingly, junctions in fiber sheaths appear flatter compared to intact filaments^8^. Thanks to the lateral resolution offered by FI-DE, we can isolate signals from the junctions. Due to the improved depth resolution, we were able to study the various profiles’ trends as a function of depth.

Profiles of the various fragments from the junctions showed similar trends (Figure 3F). Ni signal showed a subsurface peak, which comes from the conductive fibers that run parallel along with the cells and across the junction (Figure 3F, S4B). The amino acid peak is not as pronounced as that seen in the cell body. It appears that there are relatively more amino acid fragments between the two sheaths at the junction as compared to the cell body (Figure S4A). Also, the rate of a decreased intensity of the second amino acid peak from the cell junctions is lower. A look at the total number of signals of the various identified protein and carbohydrate fragments, normalized to the total counts of identified fragments, indicates that Ni’s ratio to protein fragments at the junction is lower than that of the body (Figure S5). This is in agreement with the LEXRF analysis by Boschker et al^11^. The relative amount of Ni present in the junction is identical to that of the body, suggesting that Ni is absent within the junction. Hence, the junction’s interconnects are either damaged; its composition is different from the fibers, or both. These interconnect are further studied using C-AFM.

A conductive AFM experiment was carried out on a fiber sheath by intentionally disrupting the conductive pathways identical to those performed elsewhere^10^ (Figure S5) to check whether the interconnections are present. The cuts in the filaments can be seen as trenches in the height image (Figure 5). Carbon paste, connected to the fiber sheath away from the top-left corner (not seen), acts as an electrode. The AFM probe acts as the second electrode, and wherever an electrical pathway exists between the stationary first electrode and the movable second electrode, a current flow is seen. A flow of current can be seen from the top-left until the first cut. Current flows between the first cut and the second, along the left edge. The rest of the area remains non-conductive, as they are not electrically connected to the carbon paste. This shows that the junction in the studied extracted fiber sheaths does not provide electrical interconnection of fibers, as seen in untreated filaments^10^. As already hypothesized from the FI-DE experiments, C-AFM measurements suggest that interconnects within the junction of extracted fiber sheaths are either damaged, or its composition is different from the fibers, or both. Further research is needed to resolve this issue and elucidate the cartwheel structure’s nature in the junction of cable bacteria.

## CONCLUSION

We investigated the structure and composition of cable bacteria with an enhanced lateral resolution, allowing differentiation between the cell body and the structured cartwheel junctions. The combination of ToF-SIMS FI-DE and in-situ AFM proved to be a powerful approach for this. Three ToF-SIMS modes were compared, viz., high current bunched (HCB), fast imaging (FI), and fast imaging delayed extraction (FI-DE). HCB provided the best mass resolution but lacked lateral resolution. On the other hand, FI mode provided a high lateral resolution but lacked mass resolution, which at best was in the order of a few hundreds. However, FI-DE provide a good balance between mass and lateral imaging resolution. Not only a sub-micron resolution was obtained, but the mass resolution was found sufficient to resolve signals such as Ni and other protein and carbohydrate related fragments. Ni signals reached a subsurface peak, followed by a decrease, and then reached a second maximum. This was seen now with FI-DE, but not seen earlier in HCB analysis.

Furthermore, the enhanced depth resolution was also observed by a peak in the C_4_H_8_N^+^ signal, just before the second maxima of the Ni^+^ signal. This confirms the earlier result that Ni-is present in the stillunidentified protein. The maximum intensity of CHO^+^ signal was found between the two Ni maxima, indicating a carbohydrate-rich material. By using combined ToF-SIMS/AFM, we were able to carry out both SIMS and AFM imaging in an ultra-high vacuum, avoiding further contamination when the sample is brought back into the tool after AFM imaging. Also, to find the sputtered location in an external AFM would have been difficult and time-consuming. We were able to determine the depth of subsurface maxima of Ni^+^ signal and the various depths at which carbohydrate signals were found. Comparing Ni to protein ratio in body and junction of bacteria indicated a higher ratio at the body than the junction. Also, Ni’s relative counts at the body were equal to that of the junctions, indicating Ni’s absence within the junction. This absence could be attributed to the junction and body composition differences or the interconnecting structure between adjacent cells seen in intact filaments is damaged during the extraction procedure or both. C-AFM measurements show that the interconnections at the junction seen earlier in intact filaments are not present in the fiber sheath. Not much is yet known about the composition and properties of the cartwheel structure in intact filaments. Cross-sections of intact filaments and fiber sheaths obtained by cryomicrotome and the study of these crosssections using ToF-SIMS and other complementary techniques such as C-AFM could give better insight into the composition of the cell-cell junction. From a more general analytical perspective, the proposed methodology of ToF-SIMS in FI-DE-mode, combined with AFM, could also be beneficial in resolving the structure and composition of other biological systems.

## ASSOCIATED CONTENT

### Supporting Information

The Supporting Information is available free of charge on the ACS Publications website.

The supplementary contains various ToF-SIMS analyses carried out in HCB and FI-DE mode and C-AFM setup (file type PDF) brief description (file type, i.e., PDF)

## AUTHOR INFORMATION

## Author Contributions

The manuscript was written through contributions of all authors. All authors have given approval to the final version of the manuscript

## ACKNOWLEDGMENT

The authors acknowledge Bart Cleuren and Robin Bonné for the discussion on the C-AFM experiment. The authors acknowledge the grants given by the Research Foundation – Flanders (FWO) (G031416N to RTE, JM and FJRM; G038819N to FJRM) and the Netherlands Organization for Scientific Research (NWO) (VICI grant 016.VICI.170.072 awarded to FJRM). ToF-SIMS analysis was rendered possible thanks to the grant awarded to imec vzw and Hasselt University by the Hercules foundation (now FWO; grant no. ZW/13/07 awarded to JM and AF).

## Table of contents artwork

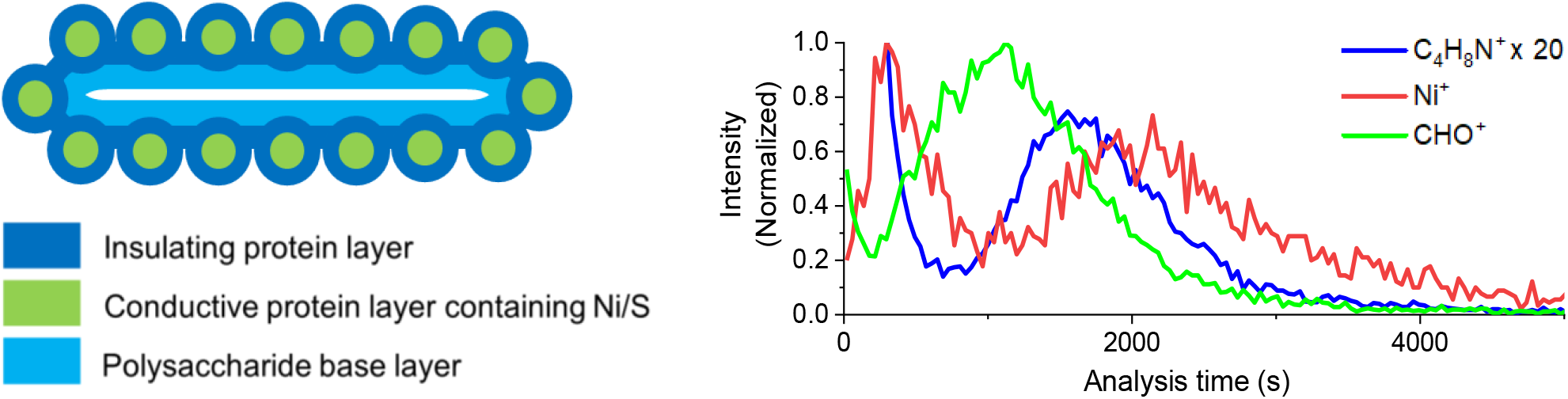

