## Supplementary information for "Enhanced laterally resolved ToF-SIMS and AFM imaging of the electrically conductive structures in cable bacteria"

**Description of figures and tables in SI**

Figure S1 is the spectra of the fiber sheath obtained in HCB mode (left) and FI-DE (right). Cycle time of 50 us was used in FI-DE, which means fragments of of m/z < 230 were collected. HCB spectra is also shown only until 230 to enable an equal comparison.

PCA analysis of fiber sheath in HCB mode is given in figure S2 A, B and in FI-DE mode in figure S2 C,D.

Figure S3 compares three typical profiles found in fiber sheath: $C_4H_8N^+$ (protein), $CHO^+$ (carbohydrate) and Ni from cell bodies and cell junctions.

Figure S4 compares the amount of all the identified protein-related and carbohydrate-related fragments and Ni in the cell body and cell junction. Signals were normalized by dividing counts of each identified fragment with the sum of counts of all identified fragments.

A schematic of the C-AFM setup is given in Figure S5.

Table S1 gives the calculated depths of $Ni^+$ and the midpoint of $CHO^+$ signal by the linear fitting of data obtained using the in-situ AFM.

Table S2 to S6 are the various fragments identified as belonging to protein layer, Nickel, carbohydrate layer, other general fragments and fragments belonging to sediments, wafer and medium. The description contains specific fragments described with 1 letter code.

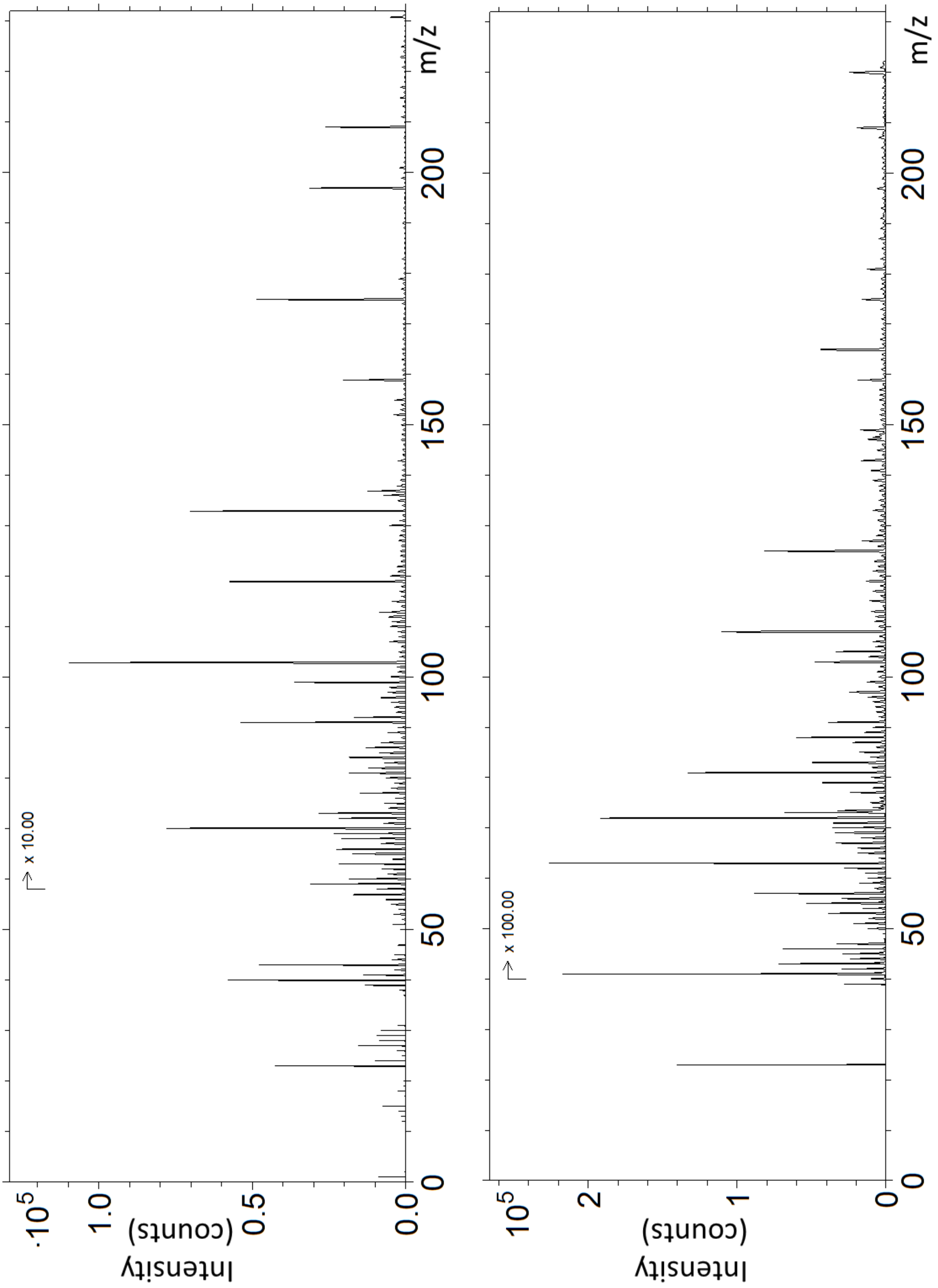


Figure S1: Spectra obtained from (left) HCB imaging and (right) FI-DE of fiber sheath.

The m/z range of HCB spectrum was chosen to reflect the available m/z range of FI DE mode due to the cycle time of 50 µs of the latter.

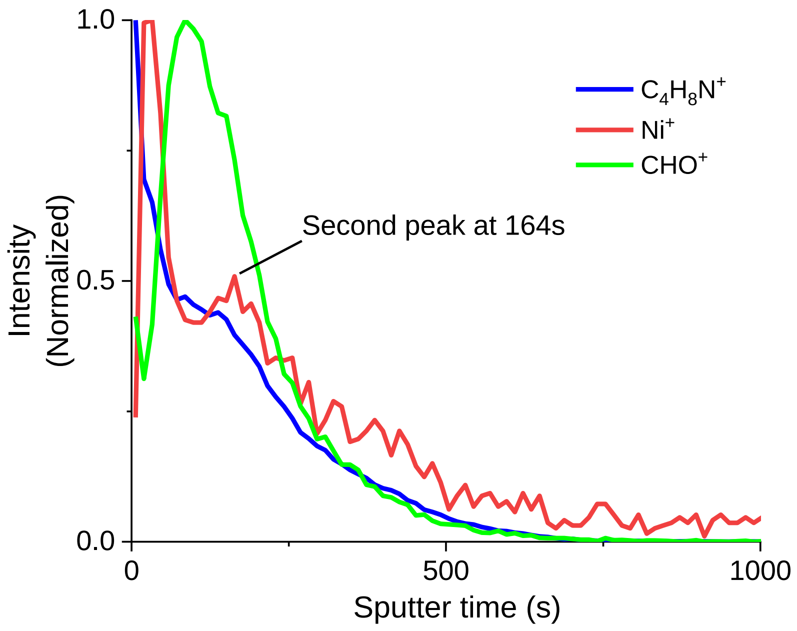


Figure S2. Trends of selected fragments as a function of sputter time in HCB mode. The first two datapoints are related to surface transient. The second peak of Ni+ is seen after 164 seconds of sputtering.

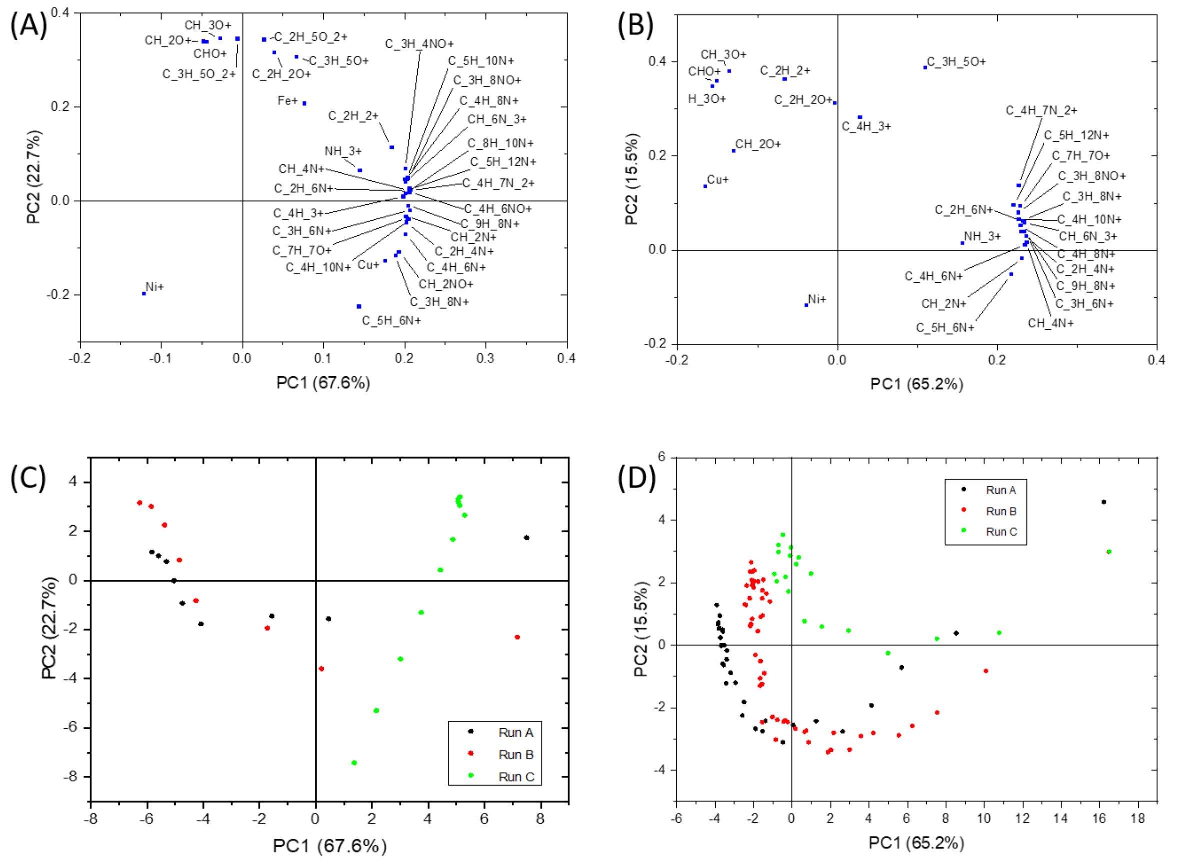


Figure S3: Principal component analysis of fiber sheath spectra obtained from HCB mode

PCA of HCB mode (A, C) and FI-DE (B, D) were obtained from three datasets each. Signals of fragments until the maximum of carbohydrate-related signals were used in PCA analysis. Amino acid fragments cluster together as are the oxygen containing carbohydrate fragments.

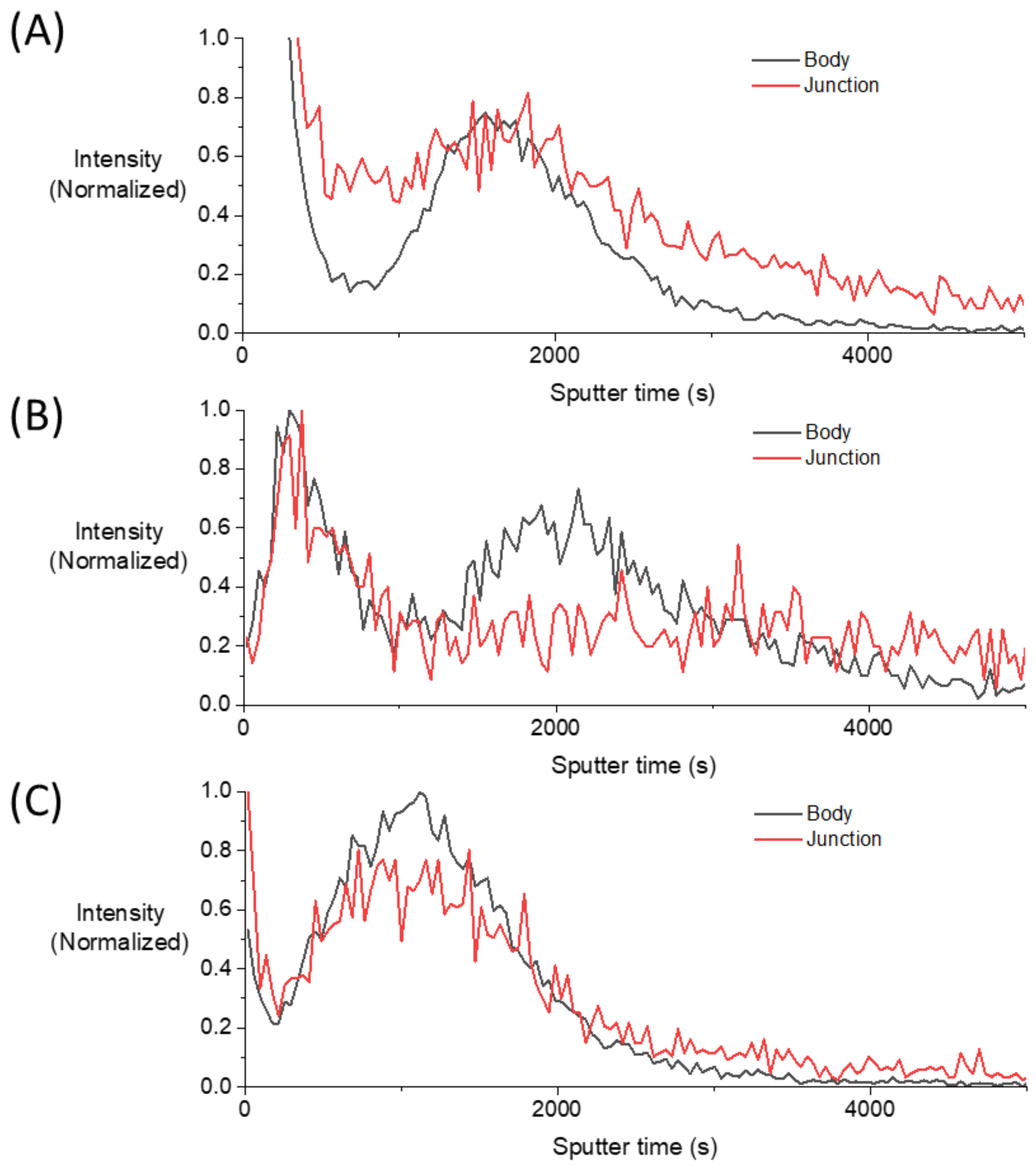


Figure S4: Comparison of three typical trends at the body and junction.

(A) Amino acid fragment $C_4H_8N^+$, (B) $Ni^+$ and (C) a carbohydrate fragment ($CHO^+$). The first peak of Ni at both body and the junction are situated at the same depth, possibly because the conductive fibers run across cells. Relative presence of amino acid fragments is higher between the top and bottom sheath at the junction compared to the body, and the rate of decrease in its intensity after the peak is slower at the junction.

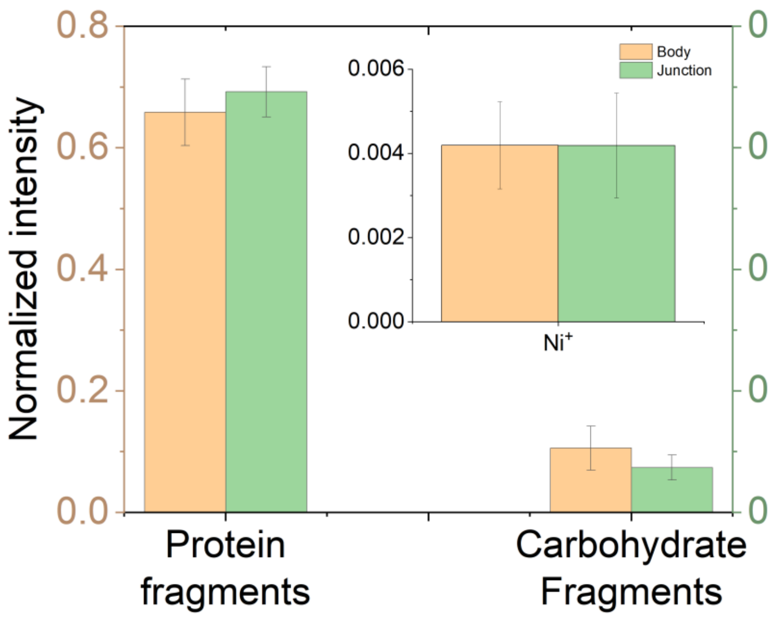


Figure S5: Comparison of protein-related, carbohydrate-related fragments and Ni. Intensities are normalized by dividing the sum of identified protein fragments, carbohydrate fragments and Ni signal with the intensity of all the identified fragments given in tables S2-S6. The error bar given is on the basis of three measurements. Cell junctions have relatively more protein fragments compared to cell body, whereas an inverse relation exists for carbohydrate fragments. Ni is identical in both parts, indicating that Ni is not present within the junction.

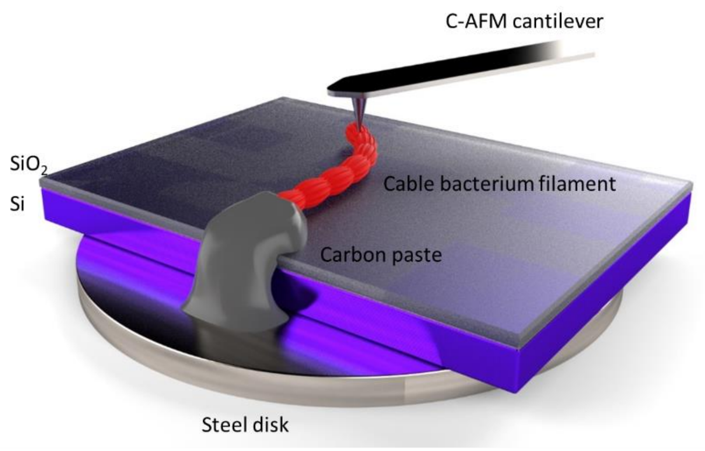


Figure S6: Schematic of conductive AFM setup.

A filament is placed over a Si wafer with 200 nm thick $SiO_2$ layer. One end of the filament is connected to the steel disk, which in turn sits on the AFM stage to which the bias voltage is applied, and the C-AFM cantilever is conductive and is connected to the ground of the controller, completing the electrical connection. When the tip sits on an exposed area of the filament, a current flows between the steel disk, through the filament to the cantilever. This is done for every pixel, resulting in the height and current images in Figure 7.

Table S1: Depth measurements of $Ni^+$ signal, $CHO^+$ signal and thickness of filament measured at the center of cell body. Depth values were obtained by a linear fit of AFM data as described in figure 6.

| Sample No. | $Ni^+$ peak (nm) | $CHO^+$ peak (nm) |
|---|---|---|
| 1 | 12.3 | 33.4 |
| 2 | 12.7 | 67.6 |
| 3 | 11.4 | 32.1 |
| 4 | 11.4 | 45.8 |
| 5 | 11.2 | 45.3 |

Table S2: Fragments associated with protein layer

| ToF-SIMS assignment | Center Mass | Expected Mass (HCB) | Expected Mass (FI - DE) | Resolution (HCB) | Resolution (FI - DE) | Description* | Used in PCA (HCB) | Used in PCA (FI - DE) |
|---|---|---|---|---|---|---|---|---|
| NH_3+ | 17.026 | 17.026 | 17.025 | 5934 | 1542 | | + | + |
| NH_4+ | 18.035 | 18.036 | 18.035 | 3557 | 1757 | | | |
| C_2H_3+ | 27.023 | 27.023 | 27.021 | 5518 | 1752 | | | |
| CH_2N+ | 28.019 | 28.019 | 28.020 | 5252 | 1538 | G | + | + |
| C_2H_4+ | 28.031 | 28.030 | | 5029 | | | | |
| CH_3N+ | 29.026 | 29.026 | | 6121 | | | | |
| C_2H_5+ | 29.040 | 29.040 | 29.039 | 5158 | 1816 | | | |
| CH_4N+ | 30.035 | 30.036 | 30.035 | 4614 | 1655 | DERMSHYFKLG | + | + |
| CH_5N+ | 31.042 | 31.042 | | 6389 | | | | |
| C_3H_5+ | 41.040 | 41.040 | 41.039 | 5406 | 1881 | | | |
| C_2H_4N+ | 42.035 | 42.035 | 42.036 | 5881 | 1745 | AF | + | + |
| CH_3N_2+ | 43.030 | 43.029 | | 7399 | | R | + | |
| C_2H_5N+ | 43.042 | 43.041 | | 6559 | | | | |
| CH_2NO+ | 44.014 | 44.012 | 44.013 | 8429 | 2622 | N | + | + |
| CH_4N_2+ | 44.037 | 44.037 | | 7481 | | R | + | |
| C_2H_6N+ | 44.052 | 44.052 | 44.049 | 5038 | 1665 | ALFYHSMKDE | + | + |
| C_4H_2+ | 50.015 | 50.015 | | 6919 | | | | |
| C_4H_3+ | 51.023 | 51.023 | 51.022 | 6446 | 2054 | | + | + |
| C_4H_5+ | 53.040 | 53.040 | 53.038 | 5859 | 2101 | | | |
| C_3H_4N+ | 54.034 | 54.035 | | 6636 | | | | |
| C_3H_3O+ | 55.020 | 55.020 | 55.013 | 5759 | 2450 | | | |
| C_3H_6N+ | 56.056 | 56.052 | 56.049 | 5231 | 1815 | FKM | + | + |
| C_3H_8N+ | 58.068 | 58.068 | 58.064 | 5302 | 1765 | E | + | + |
| C_3H_7O+ | 59.050 | 59.051 | 59.048 | 5687 | 1887 | | | |
| CH_6N_3+ | 60.058 | 60.059 | 60.054 | 5369 | 1831 | R | + | + |
| C_2H_5S+ | 61.010 | 61.012 | | 6039 | | M | | |
| C_5H_5+ | 65.040 | 65.039 | 65.036 | 6535 | 2035 | | | |
| C_4H_6N+ | 68.053 | 68.053 | | 5204 | | P | + | |
| C_4H_5O+ | 69.036 | 69.037 | | 5487 | | T | | |
| C_5H_9+ | 69.074 | 69.073 | | 5296 | | O | | |
| C_3H_4NO+ | 70.034 | 70.033 | | 5136 | | P | + | |

| | | | | | | | | |
|---|---|---|---|---|---|---|---|---|
| C_4H_8N+ | 70.071 | 70.073 | 70.065 | 4293 | 1883 | P | + | + |
| C_4H_10N+ | 72.086 | 72.088 | 72.076 | 4977 | 1847 | V | + | + |
| C_2H_7N_3+ | 73.065 | 73.065 | 73.069 | 6295 | 1738 | R | | |
| C_3H_8NO+ | 74.067 | 74.066 | 74.066 | 5217 | 1992 | T | + | + |
| C_5H_6N+ | 80.053 | 80.053 | 80.047 | 5451 | 1935 | P | + | + |
| C_4H_7N_2+ | 83.061 | 83.058 | 83.051 | 3927 | 1750 | V | + | + |
| C_4H_6NO+ | 84.048 | 84.049 | | 5344 | | EQ | + | |
| C_5H_10N+ | 84.087 | 84.090 | | 4724 | | K | + | |
| C_5H_12N+ | 86.104 | 86.104 | | 5252 | | IL | + | |
| C_4H_10N_3+ | 100.089 | 100.088 | | 5468 | | R | | |
| C_4H_8NO_2+ | 102.051 | 102.056 | | 6174 | | E | | |
| C_7H_7O+ | 107.047 | 107.054 | 107.050 | 6027 | 1923 | Y (aromatic amino acid) | + | + |
| C_5H_8N_3+ | 110.080 | 110.082 | 110.063 | 4664 | 1617 | RH | | |
| C_8H_10N+ | 120.086 | 120.087 | 120.063 | 6113 | 1721 | F (aromatic amino acid) | + | + |
| C_9H_8N+ | 130.067 | 130.068 | 130.056 | 6862 | 1948 | W (aromatic amino acid) | + | + |
| C_8H_10NO+ | 136.090 | 136.080 | 136.053 | 6029 | 1596 | | | |

Table S3: Ni signal

| ToF-SIMS assignment | Center Mass | Expected Mass (HCB) | Expected Mass (FI - DE) | Resolution (HCB) | Resolution (FI - DE) | Description | Used in PCA (HCB) | Used in PCA (FI - DE) |
|---|---|---|---|---|---|---|---|---|
| Ni+ | 57.935 | 57.934 | 57.936 | 7252 | 2073 | | + | + |

Table S4: Fragments associated with polysaccharide layer

| ToF-SIMS assignment | Center Mass | Expected Mass (HCB) | Expected Mass (FI - DE) | Resolution (HCB) | Resolution (FI - DE) | Description | Used in PCA (HCB) | Used in PCA (FI - DE) |
|---|---|---|---|---|---|---|---|---|
| O+ | 15.995 | 15.995 | 15.989 | 6070 | 1942 | | | |
| OH+ | 17.002 | 17.002 | 16.997 | 7223 | 1614 | | | |
| H_3O+ | 19.019 | 19.019 | 19.017 | 4276 | 1839 | | + | + |
| CHO+ | 29.002 | 29.002 | 29.000 | 5580 | 2087 | | + | + |
| CH_2O+ | 30.010 | 30.010 | 30.005 | 6145 | 3098 | | + | + |
| CH_3O+ | 31.021 | 31.019 | 31.016 | 5418 | 2106 | | + | + |
| C_2H_2O+ | 42.010 | 42.010 | 42.003 | 6902 | 3305 | | + | + |
| C_3H_5O+ | 45.035 | 57.036 | 57.034 | 6300 | 2463 | | + | + |
| C_2H_5O_2+ | 61.031 | 61.031 | | 6476 | | | + | |
| C_3H_3O_2+ | 71.010 | 73.031 | 71.005 | 7071 | 2953 | | + | + |
| C_3H_5O_2+ | 73.030 | 71.016 | | 6287 | | | + | |

| | | | | | |
|---|---|---|---|---|---|
| CHO_2Na_2+ | 90.980 | 90.986 | 90.973 | 3034 | 2364 |
| C_4H_2O_2Na+ | 104.999 | 104.998 | | 3860 | |

Table S5: Other general fiber sheath fragments

| ToF-SIMS assignment | Center Mass | Expected Mass (HCB) | Expected Mass (FI - DE) | Resolution (HCB) | Resolution (FI - DE) | Description | Used in PCA (HCB) | Used in PCA (FI - DE) |
|---|---|---|---|---|---|---|---|---|
| H+ | 1.007 | 1.007 | | 1901 | | | | |
| C+ | 12.000 | 11.999 | 11.997 | 5167 | 1622 | | | |
| CH+ | 13.007 | 13.007 | 13.004 | 6237 | 1474 | | | |
| N+ | 14.003 | 14.002 | | 4137 | | | | |
| CH_2+ | 14.015 | 14.015 | 13.004 | 6178 | 1474 | | | |
| NH+ | 15.010 | 15.010 | | 12287 | | | | |
| CH_3+ | 15.023 | 15.024 | 15.023 | 4493 | 1665 | | | |
| NH_2+ | 16.018 | 16.018 | 16.024 | 5650 | 981 | | | |
| C_2H_2+ | 26.015 | 26.014 | 26.011 | 5950 | 1652 | | + | + |
| CHN+ | 27.010 | 27.010 | | 7510 | | | | |
| P+ | 30.974 | 30.973 | 30.964 | 5779 | 813 | Poly-P, DNA, RNA | | |
| C_3H_2+ | 38.015 | 38.015 | 38.013 | 6914 | 1705 | | | |
| C_2H_3O+ | 43.019 | 43.019 | | 5501 | | | | |
| C_3H_7+ | 43.056 | 43.055 | 43.055 | 5537 | 2380 | | | |
| CHS+ | 44.980 | 44.979 | 44.984 | 7218 | 1912 | C | | |
| C_2H_5O+ | 45.035 | 45.035 | 45.039 | 6103 | 1713 | | | |
| PO+ | 46.969 | 46.968 | | 6815 | | Poly-P, DNA, RNA | | |
| C_3HO+ | 53.002 | 53.003 | 52.997 | 6562 | 2510 | | | |
| C_4H_7+ | 55.057 | 55.056 | 55.054 | 5462 | 1823 | | | |
| Fe+ | 55.935 | 55.934 | | 7458 | | $^{54}$Fe, $^{56}$Fe | + | |
| C_4H_9+ | 57.071 | 57.071 | 57.077 | 4840 | 2939 | | | |
| C_2H_3S+ | 58.990 | 58.995 | 59.001 | 8191 | 3131 | C | | |
| Cu+ | 62.929 | 62.929 | 62.922 | 8395 | 4696 | $^{63}$Cu, $^{65}$Cu | + | + |
| CH_2N_2O_2+ | 74.013 | 74.015 | 74.011 | 6330 | 1839 | | | |
| C_6H_3+ | 75.024 | 75.025 | 75.022 | 6227 | 2132 | | | |
| C_6H_5+ | 77.040 | 77.038 | 77.036 | 6444 | 1981 | | | |
| C_6H_6+ | 78.043 | 78.042 | 78.038 | 4137 | 2047 | | | |
| C_3H_7N_2O+ | 87.050 | 87.053 | | 4764 | | N | | |
| C_3H_6NO_2+ | 88.016 | 88.047 | | 5521 | | D | | |
| C_7H_7+ | 91.055 | 91.055 | 91.051 | 6277 | 2031 | | | |
| C_4H_4NO_2+ | 98.022 | 98.029 | | 5629 | | N | | |
| C_4H_7N_2O_2+ | 115.053 | 115.052 | 115.047 | 5486 | 1963 | G | | |

Table S6: Fragments likely derived from medium, wafer and sediment matrix

| ToF-SIMS assignment | Center Mass | Expected Mass (HCB) | Expected Mass (FI - DE) | Resolution (HCB) | Resolution (FI - DE) | Description | Used in PCA (HCB) | Used in PCA (FI - DE) |
|---|---|---|---|---|---|---|---|---|
| Na+ | 22.990 | 22.990 | 22.992 | 5216 | 1594 | Salt | | |
| Mg+ | 23.984 | 23.984 | 23.984 | 6281 | 1459 | Salt, Sediment mineral | | |
| Al+ | 26.982 | 26.981 | 26.981 | 6740 | 1604 | Sediment mineral | | |
| Si+ | 27.977 | 27.976 | 27.973 | 6651 | 1049 | Sediment mineral | | |
| K+ | 38.966 | 38.965 | 38.968 | 5344 | 1863 | Salt | | |
| Ca+ | 39.962 | 39.961 | 39.963 | 6814 | 1678 | Salt, Sediment mineral | | |
| NaCl+ | 57.958 | 57.954 | | 11936 | | Salt | | |
| KNa+ | 61.953 | 61.952 | | 4007 | | Salt | | |
| Na_2O+ | 61.975 | 61.978 | | 4569 | | Salt | | |
| Na_2OH+ | 62.985 | 62.985 | | 4737 | | Salt | | |
| NaO_3H+ | 71.985 | 71.984 | 71.986 | 6426 | 1844 | Salt | | |
| KNaOH+ | 78.959 | 78.954 | 78.952 | 3869 | 2272 | Salt | | |
| Na_2Cl+ | 80.952 | 80.963 | | 7416 | | Salt | | |
| K_2NaSO_4+ | 196.869 | 196.891 | | 6328 | | Salt | | |
| Au+ | 196.968 | 196.966 | 196.934 | 8734 | 1010 | Wafer | | |
| Au_3+ | 590.901 | 590.890 | | 10176 | | Wafer | | |
